## Supplementary material for "Nuclear VANGL2 Inhibits Lactogenic Differentiation": Rubio_Supplementary data

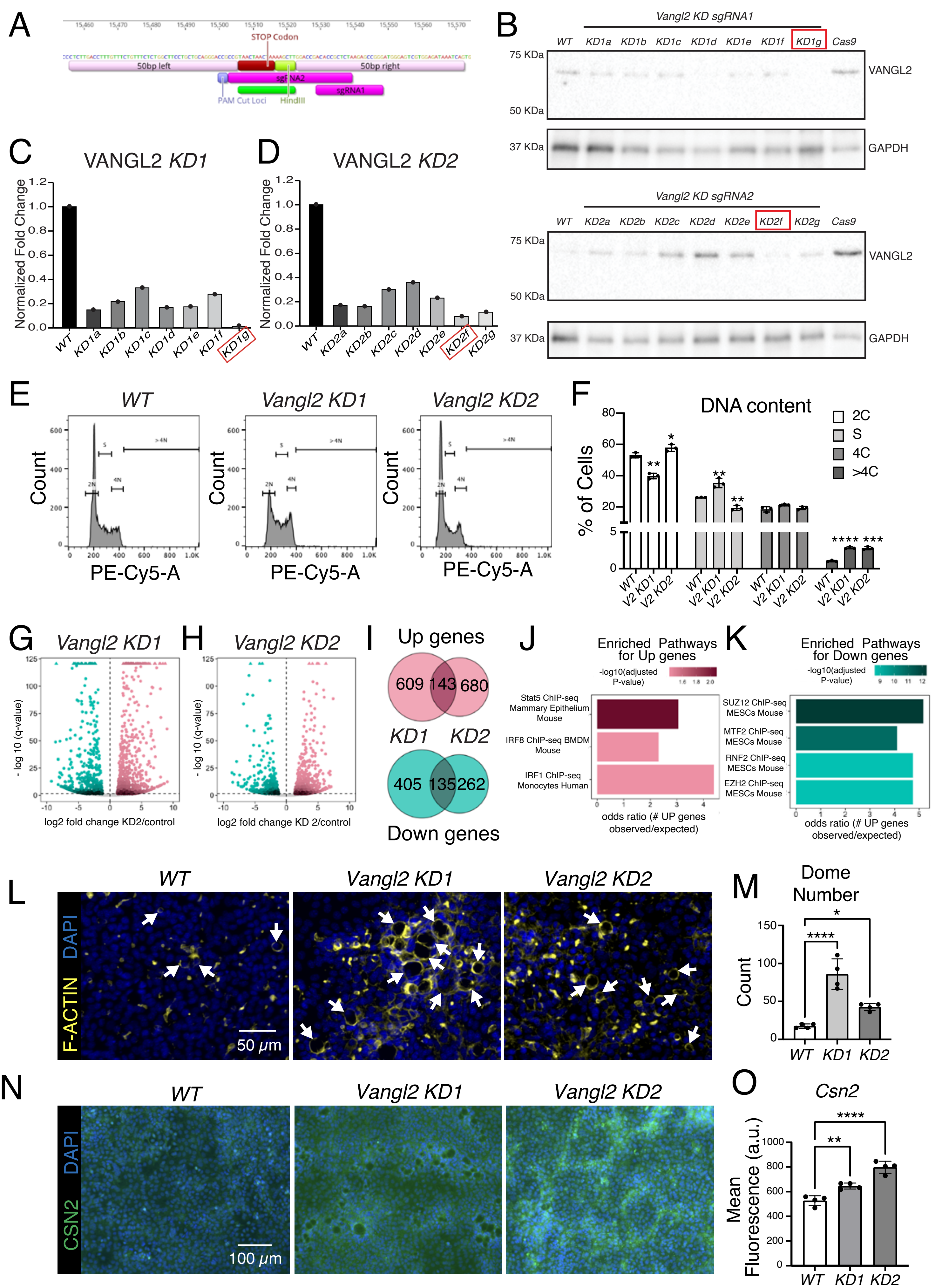

A
STOP Codon
50bp left
50bp right
sgRNA2
sgRNA1
PAM Cut Loci
HindIII
B
Vangl2 KD sgRNA1
WT KD1a KD1b KD1c KD1d KD1e KD1f KD1g Cas9
75 KDa
50 KDa
37 KDa
VANGL2
GAPDH
Vangl2 KD sgRNA2
WT KD2a KD2b KD2c KD2d KD2e KD2f KD2g Cas9
C
VANGL2 KD1
Normalized Fold Change
WT KD1a KD1b KD1c KD1d KD1e KD1f KD1g
D
VANGL2 KD2
WT KD2a KD2b KD2c KD2d KD2e KD2f KD2g
E
WT
Vangl2 KD1
Vangl2 KD2
Count
PE-Cy5-A
F
DNA content
% of Cells
2C
S
4C
>4C
WT V2 KD1 V2 KD2
G
Vangl2 KD1
- log 10 (q-value)
log2 fold change KD2/control
H
Vangl2 KD2
log2 fold change KD 2/control
I
Up genes
609 143 680
KD1 KD2
405 135 262
Down genes
J
Enriched Pathways for Up genes
-log10(adjusted P-value)
Stat5 ChIP-seq Mammary Epithelium Mouse
IRF8 ChIP-seq BMDM Mouse
IRF1 ChIP-seq Monocytes Human
odds ratio (# UP genes observed/expected)
K
Enriched Pathways for Down genes
SUZ12 ChIP-seq MESCs Mouse
MTF2 ChIP-seq MESCs Mouse
RNF2 ChIP-seq MESCs Mouse
EZH2 ChIP-seq MESCs Mouse
L
WT
Vangl2 KD1
Vangl2 KD2
DAPI
F-ACTIN
50 μm
M
Dome Number
Count
WT KD1 KD2
N
WT
Vangl2 KD1
Vangl2 KD2
DAPI
CSN2
100 μm
O
Csn2
Mean Fluorescence (a.u.)
WT KD1 KD2

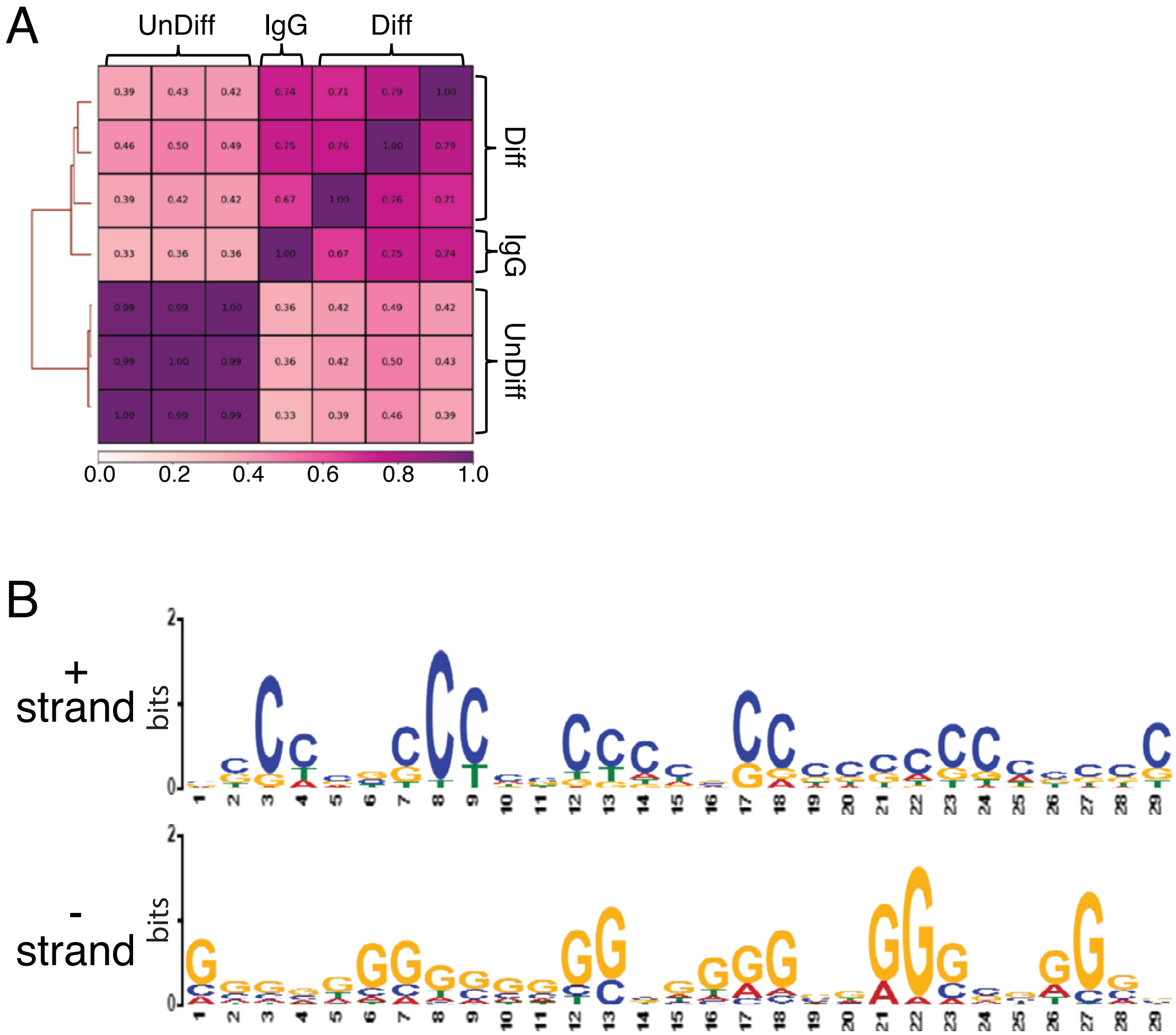
A
UnDiff
IgG
Diff
0.0
0.2
0.4
0.6
0.8
1.0
B
+
strand
bits
-
strand
bits

### SUPPLEMENTARY FIGURE LEGENDS

**Supplementary Figure 1. *Vangl2* KO cells have express proteins and genes involved in lactogenic differentiation.**

(A) Cartoon representation of CRISPR/Cas9 strategy to knockdown *Vangl2* using two different RNA guide sets.

(B-D) Western blots (B) and quantification of VANGL2 in *Vangl2 KD*1 (C) and *Vangl2 KD* 2 (D) HC11 cells. Red boxes indicate *KD1* and *KD2* HC11 cells selected for further study.

(E) Representative FACS histograms for DNA content analysis of undifferentiated *WT*, *Vangl2 KD1* and *Vangl2 KD2* HC11 cell lines as labeled by propidium iodide (PE-Cy5-A).

(F) Quantification of FACS DNA content analysis showing the percentage of undifferentiated *WT*, *Vangl2 KD1* and *Vangl2 KD2* HC11 cells with 2C, 4C or > 4C DNA content or in the S phase.

(G, H) Volcano plots showing the differential gene expression of the *Vangl2 KD1* and *Vangl2 KD2* cells in comparison to *WT*. Each dot represents a gene. Teal color shows genes with decreased expression and pink shows genes with increased expression.

(I) (Top) Venn diagram showing the number of upregulated (Up) genes in *Vangl2 KD1* (609) and in *Vangl2 KD2* (680). (Bottom) Venn diagram showing the number of downregulated (Down) genes in *Vangl2 KD1* (405) and *Vangl2 KD2* (262). The shared differentially expressed genes (Up: 143, Down: 135) were used to generate the enriched pathway plots.

(J, K) Enriched pathways from the genes upregulated in *Vangl2 KD1* cells (J) and downregulated in *Vangl2 KD2* cells (K). The colored scale is logarithmic, with, the darker colors representing higher p-values.

(L, M) Representative immunofluorescence microimages of *WT*, *Vangl2 KD1* and *Vangl2 KD2* HC11 cells at differentiation day 2 immunostained for F-actin (yellow), with nuclei labeled with Hoechst (blue) L), and quantification of the number of domes (arrows) (M).

(N, O) Representative images of *WT*, *Vangl2 KD1* and *Vangl2 KD2* HC11 cells at differentiation day 2 immunostained for CSN2 (green), with nuclei labeled with Hoechst (blue) (N), and quantification of CSN2 mean fluorescence (O).

Error bars represent mean ± SD. Error bars represent mean ± SD. Analyzed by two-tailed un-paired Student's t-test. n=3 biological replicates. p values: * < 0.05, **< 0.01, *** < 0.001, **** < 0.0001.

**Supplementary Figure 2. VANGL2 binds DNA motifs in undifferentiated, but not differentiated, HC11 cells, including one in the *Stat5a* promoter.**

(A) Heatmap of the Pearson correlation measure over the genome-wide signal for CUT&RUN reads of undifferentiated (unDiff) and differentiated (Diff) HC11 cells using a rabbit anti-VAGNLL2 antibody or non-specific rabbit IgG. Rows are clustered according to inter-sample correlation. Color intensity is proportional to the correlation value reported in each cell.

(B) Representation of the motif found in the *Stat5a* promoter (forward + and reverse – DNA strand).
